# *In vitro* trypanocidal effect of sera from *Erythrocebus patas* (Red Patas monkey) and *Chlorocebus tantalus* (Tantalus monkey) on *Trypanosoma brucei brucei* Plimmer & Bradford, 1899 and *Trypanosoma congolense* Broden, 1894

**DOI:** 10.1101/2020.07.13.200311

**Authors:** Felicite Djieyep-Djemna, Ishaya Haruna Nock, Thelma Aken’Ova, Ezekiel Kogi, Armand Claude Noundo Djieyep

## Abstract

Anti-*Trypanosoma brucei brucei* and anti-*Trypanosoma congolense* activities of sera from two species of uninfected zoo-primates, *Erythrocebus patas* (red patas monkey) and *Chlorocebus tantalus* (tantalus monkey) were investigated. The sera were screened using thick films and haematocrit centrifugation technique (HCT), to ensure that the donor primates were not infected with trypanosomes. *Trypanosoma brucei brucei* (Federe strain) and *Trypanosoma congolense* were suspended in supplemented RPMI (Rossvelt Park Memorial Institute) 1640 medium and the motility of the parasite was used as index of viability after the addition of each test serum. The selected primate sera exhibited some degree of anti-*Trypanosoma brucei brucei* activities *in vitro*. Red patas monkey serum had an inhibition index of 0.27, while that of Tantalus monkey was 0.34, against *Trypanosoma brucei brucei*, with mean survival times of 22.00±1.73 hours for red patas monkey serum and 19.67±0.58 hours for tantalus monkey serum, which are significantly lower (P<0.05) than that of the control (30.00±0.00 hours). The selected primate sera had pronounced inhibitory activities against *Trypanosoma congolense*. Sera from the two species of primate had very high anti-*Trypanosoma congolense* activity showing an inhibition index of 0.91 for Red patas monkey serum and 0.90 for Tantalus monkey serum, with marked and significant reduction (P<0.05) in survival time of 7.00±1.73 hours in Red patas monkey serum and 7.67±0.58 hours in Tantalus monkey serum, compared with the control (74.00±1.00 hours). The *in vitro* anti-trypanosomal activity of the serum samples was shown to be cidal in nature. The activity was not associated with xanthine oxidase. This study revealed that sera from red patas monkey and tantalus monkey had a moderate anti-*Trypanosoma brucei brucei* activity and a very high anti-*Trypanosoma congolense* activity *in vitro* suggesting the presence of some non-specific materials.

**Authors’ Summary:** The mechanisms that allow trypanosomiasis-resistant animals to control blood trypanosomes are being investigated, to identify non-specific factors that kill trypanosomes or limit their proliferation, contributing to host resistance. For instance, xanthine oxidase has been isolated and identified as the protein that kills trypanosomes in Cape buffalo. Humans and several other primates are also known to be resistant to infection by several animal-specific trypanosome species. In this study, sera from some zoo primates, red patas monkey and tantalus monkey, tested on *Trypanosoma brucei brucei* and *Trypanosoma congolense in vitro*, showed a slight anti-*Trypanosoma brucei brucei* activity and a very high anti-*Trypanosoma congolense* activity. These activities were shown to be cidal in nature and not associated with the protein xanthine oxidase. The authors suggest that non-specific factors other than the enzyme xanthine oxidase might have accounted for the sera anti-trypanosomal activities.

## Introduction

African animal trypanosomiasis constitutes a major impediment to efficient and profitable livestock production in Africa generally and particularly in Nigeria. It causes about 3 million deaths annually in cattle and production losses of about US$ 1.2 billion [1]. Although several measures have been used to attempt the control of the disease, the main control measure remains the use of trypanocidal drugs since the antigenic variation in the parasite has render the production of vaccine (using antibody) a very difficult task. Thus trypanosomiasis is still a great challenge to scientists. New control approaches are based on nonspecific factors. Some wild animals are known to be naturally trypanosomiasis-resistant. Natural resistance is attributable to natural specific antibodies and nonspecific serum factors capable of killing trypanosomes or limiting their rate of population growth, thereby contributing to the trypanotolerance/trypanoresistance of some animals. For instance, 3 distinct constitutive defense mechanisms against *T. brucei* have been detected by Black *et al*. [2], in sera from sub-Saharan mammals other than primates that show a high level of resistance to African trypanosomes, and they reported that these mechanisms act together with parasite-specific antibody responses to control the severity of infection arising in the reservoir hosts [2]. In the same regard, some studies have also been carried out on some zoo primates [3], [4], [5]. The present study was targeting at investigating the presence of nonspecific factors against *T. brucei brucei* and *T. congolense*, in the sera of two species of Zoo Primates, *Erythrocebus patas* (Red patas monkey) and *Chlorocebus tantalus* (Tantalus monkey).

## Materials and methods

### Ethical statement

The ethical approval for this research was obtained from the Committee on Animal Use and Care, Directorate of Academic Planning & Monitoring, Ahmadu Bello University, Zaria, with the Approval No: ABUCAUC/2017/007. Local approval was given by the Director of Kano Zoological Garden. Blood samples were collected in accordance with best practice guidelines to minimise stress on donor primates. The laboratory animals used were kept in a clean aerated room and were given necessary care.

### Donnor primates and sampling sites

Blood was collected from red patas monkey and tantalus monkey reared in an area free of trypanosomes, the Kano Zoological Garden, Kano State, Nigeria. Kano is 481 metres (1,578 feet) above sea level. The city lies to the north of the Jos Plateau, in the Sudanian Savanna region that stretches across the south of the Sahel. The city lies near where the Kano and Challawa rivers flowing from the southwest converge to form the Hadejia River, which eventually flows into Lake Chad to the east [6].

### Sample collection and trypanosome screening test

This was done with the assistance of some experts. Each blood sample was aseptically collected from the donor animal through the femoral vein and was divided into two. One part was kept as whole using EDTA (ethyl-diamine-tetra-acetic acid) while the other one was dispensed into sterile plain glass test tube, allowed to clot for 2-3 hours at room temperature and serum was collected after centrifugation at 1,000 g [2] and stored in a freezer at -20°C after screening, for further analysis. The screening was done on each of the blood sample with EDTA, using two of the standard detection techniques (thick films) and haematocrit centrifugation technique (HCT) [7].

### In vitro detection of anti-trypanosomal activity of the test sera

This was done using microtitre plates as follows:

- Trypanosomes from the buffy coat were suspended in RPMI 1640 (Rossvelt Park Memorial Institute 1640) medium supplemented with 2% glucose, 2mM sodium pyruvate, 10% heat-inactivated (56 C, 30 min) fetal bovine serum [8], sodium bicarbonate and sodium pyruvate and antibiotics (streptomycin 100µg/ml, penicillin 100U/ml).
- 50µl of each monkey serum sample was introduced into one of the 96 wells of a microtitre plate and 50µl of *T. brucei brucei* or *T. congolense* suspension 8 per field (31.62×10^6^/ ml of blood) was added to it, rocked gently to mix and incubated at room temperature.
- A drop of about 5µl of each mixture was examined microscopically, hourly using wet film method. Cessation in motility of parasites was taken as indication of serum activity against the parasites [9].
- The motility of *T. brucei brucei* or *T. congolense* in each well was compared with the motility of the same parasites in the control well without test serum.

A formula was derived [10] to determine the Anti-trypanosomal Activity Index (ATI) of each serum sample.

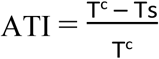

Where: T_c_ is the survival time of the parasites in the control medium

T_s_ is the survival time of the parasites in the sample

### Infectivity assessment

100µl of the mixture of the *in vitro* affected parasite with effective serum was inoculated into mice and monitored by microscopy for trypanosomes on daily basis, 10 and 30 days for *T. brucei brucei* and *T. congolense* respectively, to determine if the observed anti-trypanosomal activity was inhibitory or cidal.

### Detection of the xanthine oxidase content of the selected sera

The technique of Black *et al*. [2] was adopted to detect the xanthine oxidase content of the test sera. Accordingly, aliquots (100 µl) of each serum was added to 900 µl of H_2_0_2_-assay buffer (0.5 mM xanthine and 1 mM 2, 4, 6 tribromo-3-hydroxybenzoic acid in 0.1 mM 4-amino-antipyrine with a final concentration of 8 units horse-radish peroxidase per ml). The mixture was incubated at 25°C for 30 min, immediately chilled in an ice bath, and absorbance read at 512 nm wave length was recorded after zeroing the spectrophotometer with an equivalent mixture lacking horseradish peroxidase. A serial dilution of the commercial cow’s milk xanthine oxidase was done; absorbance was also recorded at 512 nm wave length and used to plot a standard curve. It has been established that detection of H_2_0_2_ produced in serum by this assay is not affected by other enzymes in serum, including catalase [11]. The xanthine oxidase content of serum was determined by reading the serum value against the cow’s milk xanthine oxidase standard curve.

### Data analysis

One-way Analysis of variance (ANOVA) was used for sera’s inter-species comparison of the parasites survival time. Student’s t-test was used to compare the mean survival time of the two species of parasites, *T. brucei brucei* and *T. congolense* in the test sera. The significant difference was at the level of probability 0.5. All data were expressed as means ± Standard Error.

## Results

### Trypanosome Infection Status of Sera from Selected Primates

Blood samples from selected monkeys were negative for trypanosomes by thick blood film and haematocrit centrifugation technique (HCT).

### The effect of sera from Red patas monkey and Tantalus monkey on T. brucei brucei

There was no significant difference in the trypanosomal activity of the sera of both primates (22.00±1.73 and19.67±0.58 hours respectively); meanwhile, the mean survival times of the parasites in these sera were significantly lower (P<0.05) than that of the control, the RPMI 1640 (30.00±0.00 hours), showing a slight anti-*Trypanosoma brucei brucei* activity with ATI values of 0.27 and 0.34 respectively (Table 1).

**Table 1:** Effect of sera of selected Zoo Primates on *T. brucei brucei in vitro*.

| Source of serum | Survival time (hrs) | ATI |
| --- | --- | --- |
| *Erythrocebus patas* (Red Pattas monkey) | 22.00±1.73<sup>a</sup> | 0.27 |
| *Chlorocebus tantalus* (Tantalus monkey) | 19.67±0.58<sup>a</sup> | 0.34 |
| RPMI 1640 | 30.00±0.00<sup>b</sup> | 0 |

Values are mean ± standard deviation. Values with different superscripts down the column are significantly different (P<0.05). ATI: Anti-trypanosomal activity Index.

### The effect of sera from Red patas monkey and Tantalus monkey on T. congolense

Sera from the selected zoo primates exhibited very high anti-*Trypanosoma congolense* activity *in vitro* with very high index, close to one (ATI of 0.91 for Red pattas monkey and 0.90 for Tantalus monkey). This was shown by a very reduced survival time of *T. congolense* 7.00±1.73 hours in Red pattas monkey serum and 7.67±0.58 hours in Tantalus monkey (Table 2). These two values are not statistically different (P>0.05) from each other, but both are significantly (P<0.05) lower than the control, the RPMI 1640 (74.00±1.00 hours).

**Table 2:** Effect of sera of selected Zoo Primates on *T. congolense*.

| Source of serum | Survival time (hrs) | ATI |
| --- | --- | --- |
| *Erythrocebus patas* (Red Pattas monkey) | 7.00±1.73<sup>a</sup> | 0.91 |
| *Chlorocebus tantalus* (Tantalus monkey) | 7.67±0.58<sup>a</sup> | 0.90 |
| **RPMI 1640** | 74.67±1.00<sup>b</sup> | 0 |

Values are mean ± standard deviation. Values with different superscripts down the column are significantly different (P<0.05). ATI: Anti-trypanosomal activity Index.

### The compared effect of sera from Red patas monkey and Tantalus monkey on T. bucei brucei and T. congolense in vitro

Each of these sera had a very high anti-*Trypanosoma congolense* activity Index (ATI), 0.91 for Red pattas monkey and 0.90 for Tantalus monkey. This was very high compared to their activity index against *T. brucei brucei* (0.27 for Red patas monkey and 0.34 for Tantalus monkey). The survival times of *T. congolense* in these sera are significantly lower (P<0.05) than that of *T. brucei brucei* in the same sera (Table 3).

**Table 3:** Compared effect of sera from selected Zoo primates on *T. brucei brucei* and *T. congolense in vitro*.

| Source of serum | Survival<br>time of *T.*<br>*brucei* | ATI<br>of the serum<br>on *T.*<br>*brucei* | Survival<br>time of *T.*<br>*congolense* | ATI of the<br>serum on<br>*T.*<br>*congolense* |
| --- | --- | --- | --- | --- |
| **Red Pattas monkey (*Erythrocebus patas*)** | 22.00±1.73<sup>b</sup> | 0.27 | 7.00±1.73<sup>a</sup> | 0.91 |
| ***Chlorocebus tantalus* (Tantalus monkey)** | 19.67±0.58<sup>b</sup> | 0.34 | 7.67±0.58<sup>a</sup> | 0.90 |
| **RPMI 1640** | 30.00±0.00<sup>a</sup> | 0 | 74.00±1.00<sup>b</sup> | 0 |

Values are mean ± standard deviation. Values with different superscripts across the rows are significantly different ( $P < 0.05$ ). ATI: Anti-trypanosomal activity Index.

### Infectivity

None of the parasites affected by the test sera *in vitro* was able to cause infection to the mice.

### Xanthine oxidase content of sera of selected zoo primates

The xanthine oxidase content of sera of the selected zoo primates is presented in Figure 1. It was observed that concentration of xanthine oxidase in the serum of red patas monkey was significantly higher (P<0.05) than that of Tantalus monkey (2.26±0.10 and 1.02±0.33 µg/ml respective.

**Figure 1:**
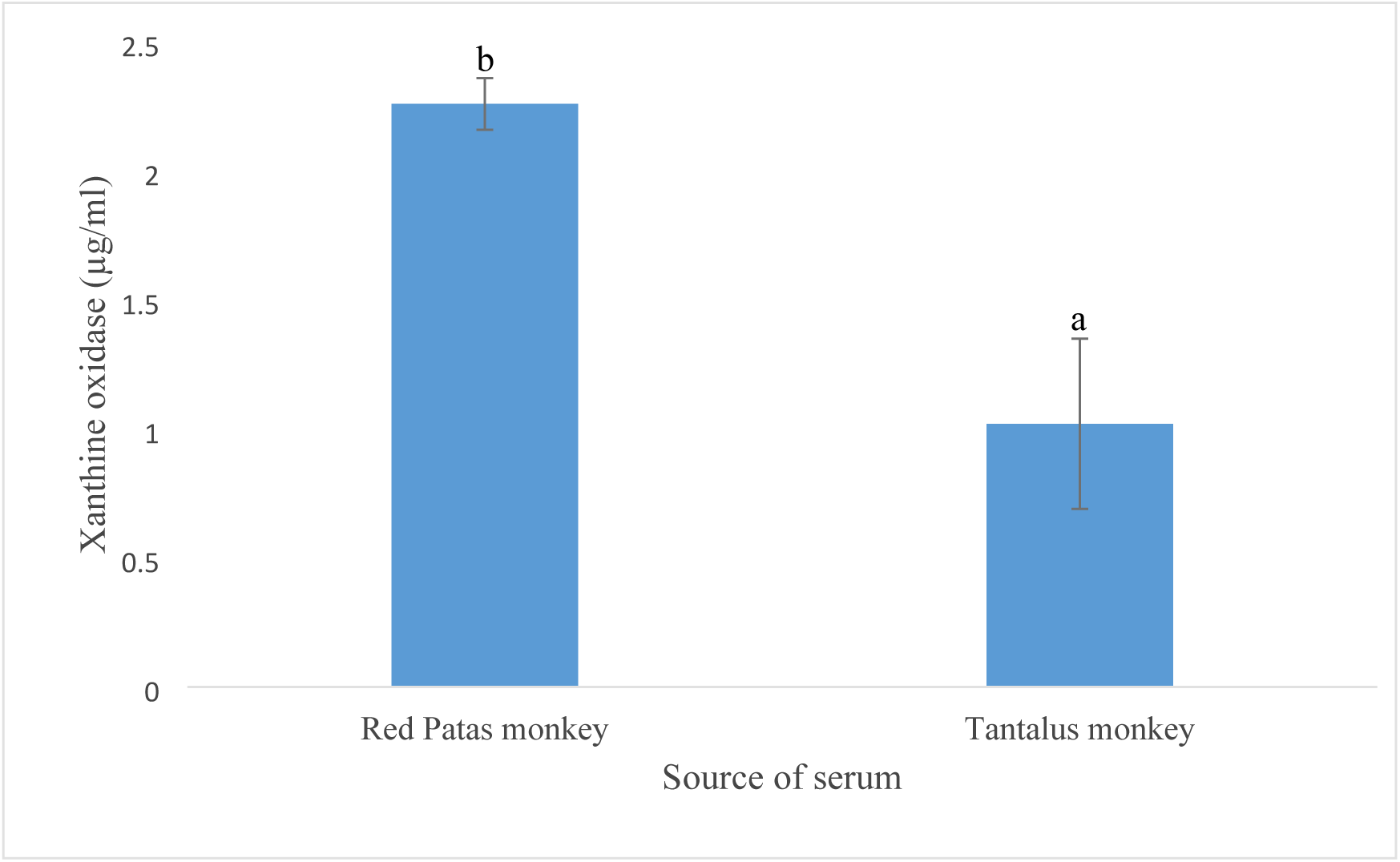
Xanthine oxidase content of sera of selected zoo primates.

Values are mean concentrations. Values with different superscripts are significantly different (P<0.05).

## Discussion

A simple technique was employed using the motility of trypanosomes as indicator of parasites viability [9] as it has long been established that parasites motility could be a measure of viability among most zooflagelate parasites [12]. Also, Atawodi *et al*. [13] reported that the technique correlated well with other *in vitro* methods.

The fact that sera from the two zoo primates, Red patas monkey and Tantalus monkey showed a slight anti-*Trypanosoma brucei brucei* activity Index (ATI of 0.27 and 0.34 respectively) and a very high anti-*Trypanosoma congolense* activity Index *in vitro* (ATI of 0.91 and 0.90 respectively) with a highly reduced mean survival time of the parasites of about 7.00±1.73 hours for red patas monkey and 7.67±0.58 hours for tantalus monkey, demonstrate an innate immunity of these primates to *T. congolense*, and to some lesser extend to *T. brucei brucei*. Human and several other primates are known to be resistant to infection by several animal-specific trypanosome species including *T. brucei brucei, T. congolense, T. vivax* and *T. evansi* [3]. The results obtained suggest that Red pattas monkey serum as well as Tantalus monkey serum might possess some trypanolytic factors against *T. congolense*, and to some lesser extend against *T. brucei brucei*. This claim can be supported by the fact that the innate immunity of human and several other primates to the trypanosomes is due to the expression of a unique subset of high-density lipoproteins (HDLs) referred to as the trypanosome lytic factors (TLFs) in their blood [3]. Being that the trypanosome lytic factor is primarily composed of Apolipoprotein L1 (APOL1) and a haptoglobin-related protein [14], and considering the findings of Jirku *et al*., [4] who demonstrated that mandrill serum was able to efficiently lyse *T. brucei brucei* and *T. brucei rhodesiense*, and to some extent T. *bucei gambiense*, while the chimpanzee serum failed to lyse any of these subspecies because of the secondary loss of the APOL1 gene, the anti-trypanosomal activity of red pattas monkey and tantalus monkey sera observed in this study could be attributable to Apolipoprotein L1 (APOL1). These findings also correlate with some previous ones among which the identification of an Apolipoprotein L1 (APOL1) in a subset of Old World monkeys [15], [16], [17], [18], the demonstration of the *in vitro* lytic ability of serum and purified recombinant protein of an Apolipoprotein L1 (APOL1) ortholog from the West African Guinea baboon (*Papio papio*), which is able to lyse all subspecies of *T. brucei* including *T. brucei gambiense* [5].

The parasites were inactivated by the monkeys’ sera *in vitro* and unable to cause infection to mice suggesting that the anti-trypanosomal activity observed could be cidal. Considering the fact that the sera were obtained from primates that had not previously been exposed to trypanosome infection, these results are also suggestive of the nonspecific nature of the anti-trypanosomal materials present in the sera. The difference in the xanthine oxidase content of sera from red patas monkey and Tantalus monkey indicates that the similar anti-trypanosomal activity observed is not associated to the activity of this enzyme. This finding gives more support to our suggestion on the possible trypanolytic factor previously mentioned.

## Conclusion

This study revealed anti-*Trypanosoma brucei brucei* and anti-*Trypanosoma congolense* properties of the sera of the two zoo primates, Red patas monkey and Tantalus monkey *in vitro*. Both sera had slight anti-T*rypanosoma brucei brucei* activity and a very high anti-T*rypanosoma congolense* activity *in vitro*. These activities, showing some innate immunity is attributable to some nonspecific factors (trypanolytic factors) present in the sera of the Red patas monkey and the Tantalus monkey.

## Acknowledgement

The authors would like to thank the officers and staff of the Kano Zoological garden, Kano state, Nigeria, for approving the collection of blood samples from their animals. We are also grateful to the managements of the Departments of Microbiology, Faculty of Life Sciences, Parasitology and Entomology Faculty of Veterinary Medicine all of the Ahmadu Bello University, Zaria, for permitting us to use their facilities and their technical staffs.

## References

(1) Chamond, N., Cosson, A., Blom-Potar, M.C., Jouvion, G., D’Archivio, S., Medina, M., Droin-Bergère, S., Huerre, M., Goyard, S., Minoprio, P. (2010). *Trypanosoma vivax* Infections: Pushing Ahead with Mouse Models for the Study of *Nagana*. I. Parasitological, Hematological and Pathological Parameters. PloS Neglected Tropical Diseases, 10;4 (8): e792.

(2) Black, S.J., Wang, Q., Makadzange, T., Li, Y., Van Praagh, A., Loomis, M. and Seed, J.R. (1999). Anti-trypanosoma brucei activity of nonprimate zoo sera. American Society of Parasitology, Journal of Parasitology 85 (1): 48–53.

(3) Namangala, B. (2011). How the African trypanosomes evade host immune killing. Parasite Immunology. 33 (8):430–7. doi: 10.1111/j.1365-3024.2011.01280x.

(4) Jirků, M., Votýpka, J., Petrželková, K.J., Jirků-Pomajbíková, K., Kriegová, E., Vodička, R., Lankester, F., Leendertz, S.A.J., Wittig, R.M., Boesch, C., Modrý, D., Ayala, F.J., Leendertz, F.H. and Lukeš, J. (2015). Wild chimpanzees are infected by *Trypanosoma brucei*. International Journal of Parasitology. Parasites Wildl, 4 (3): 277–282.

(5) Cooper, A., Capewell, P., Clucas, C., Veitch, N., Weir, W., Thomson, R., Raper, J., MacLeod, A. (2016). A Primate APOL1 Variant That Kills *Trypanosoma brucei gambiense*. PLOS Neglected Tropical Diseases, 10 (8): e0004903. doi: 10.1371/journal.pntd.0004903. eCollection 2016.

(6) Kano. Available from: https://en.wikipedia.org/wiki/Kano

(7) Woo, P.T.K. (1989), The haematocrit centrifuge for the detection of trypanosomes in blood. Canadian Journal of Zoology. 47: 921–923.

(8) Black, S.J. and Vandeweerd, V. (1989). Serum lipoproteins are required for multiplication of Trypanosoma brucei under axenic conditions. Molecular and Biochemical Parasitology, 37: 65–72.

(9) Bulus, T., Atawodi, S.E. and Mamman, M. (2008). *In vitro* Antitrypanosomal Activity and Phytochemical Screening of Aqueous and Methanol Extracts of *Terminalia avicennioides*. Nigerian Journal of Biochemistry and Molecular Biology, 23 (1): 7 – 11.

(10) Djieyep-Djemna, F., Kogi, E., Nock, I.H., Aken’Ova, T.O.L. (2017). A Model for Determination of *In Vitro* Trypanosomal Activity Status Of Sera Using An Anti-Trypanosoma Activity Index (ATI). Parasitology and Public Health Society of Nigeria. Abstract No. PPSN/2017/ABS/115. P. 84.

(11) Le Tissier, P. R., Peters, J. and Skidmore, C. J. (1994). Development of an assay method for purine catabolic enzymes in the mouse and its adaptation for use on an autoanalyzer. Analytical Biochemistry, 222:168–175.

(12) Peter, D., Honigberg, B.M. and Fern, A.M. (1976). An improved method of cryopreservation of *Trypanosoma (Nannomonas) Congolense* brooden in liquid nitrogen. Journal of Parasitology, 62 (1): 136–137.

(13) Atawodi, S.E., Bulus, T., Ibrahim, S., Ameh, D.A., Nok, A.J., Mamman, M., Galadima, M. (2003). *In vitro* trypanocidal effect of methanolic extract of some Nigerian savannah plants. African Journal of Biotechnology, 2 (9): 317–321.

(14) Raper, J., and Friedman, D.J. (2013). Parasitology: molecular one-upmanship. Nature, 501:322–323.

(15) Lugli, E.B., Pouliot, M., Portela, M.D.P.M., Loomis, M.R., Raper, J. (2004). Characterization of primate trypanosome lytic factors. Molecular and Biochemical Parasitology, 138(1):9–20. pmid:1550091.

(16) Poelvoorde, P., Vanhamme, L., Van Den Abbeele, J., Switzer, W.M., Pays, E. (2004). Distribution of apolipoprotein L-I and trypanosome lytic activity among primate sera. Molecular and Biochemical Parasitology, 134:155–157. pmid:14747153.

(17) Thomson, R., Molina-Portela, P., Mott, H., Carrington, M., Raper, J. (2009). Hydrodynamic gene delivery of baboon trypanosome lytic factor eliminates both animal and human-infective African trypanosomes. Proceeding of National Academic Sciences USA.106: 19509–19514.

(18) Thomson, R., Genovese, G., Canon, C., Kovacsics, D., Higgins, M.K, Carrington, M. (2014). Evolution of the primate trypanolytic factor APOL1. Proceedings of National Academic Sciences, USA. National Academic Sciences, 111: 2130–2139.

